# CP12 controls ribulose 1,5 bisphosphate recycling and carbon acquisition in *Chlamydomonas reinhardtii*

**DOI:** 10.1101/2025.06.13.659560

**Authors:** Cassy Gérard, Régine Lebrun, Christophe Verthuy, Hugo Le Guenno, Artemis Kosta, Florence Guérard, Kwang Suk Chang, Luisana Avilan, Bertrand Gakière, EonSeon Jin, Stephen C. Maberly, Brigitte Gontero, Hélène Launay

## Abstract

The small chloroplastic protein CP12 has multiple functions, including the regulation of enzymes in the Calvin-Benson-Bassham cycle. Here, we investigated its role in the acclimation of *Chlamydomonas reinhardtii* to varying CO_2_ availability. This alga has a CO_2_ concentrating mechanism that increases the supply of CO_2_ to ribulose-1,5-bisphosphate carboxylase/oxygenase (RuBisCO) and involves hallmarks such as HCO_3_^-^ transporters and carbonic anhydrases as well as the condensation of RuBisCO within the pyrenoid via its interaction with a scaffold protein named Essential Pyrenoid Component 1 (EPYC1). We showed that compared to the wild type, at high CO_2_, *C. reinhardtii* CP12 deletion mutants, or partially complemented mutants, have less phosphoribulokinase and ribulose-1,5-bisphosphate (RuBP) indicating that the regeneration of RuBP is regulated by CP12. In the absence of CP12, the expected relocation of RuBisCO towards the pyrenoid was not observed upon transition from high to very low CO_2_, contrary to WT cells. The CP12 deletion mutants are a unique example where the induction of CO_2_ concentrating mechanism hallmarks at very low CO_2_ was not accompanied by RuBisCO relocation. Altogether, these results suggest that CP12 contributes to the coordination between RuBP regeneration, RuBisCO location and CO_2_ acquisition.

**Highlight:** CP12 regulates phosphoribulokinase amount and its product RuBP. The CP12 deletion mutants are unique example where the RuBisCO location was not correlated with the induction of CO_2_ concentration mechanism. This reveals a novel link between Calvin-Benson-Bassham cycle regulation and CO_2_ concentrating mechanisms in *Chlamydomonas reinhardtii*.

**Graphical abstract:** 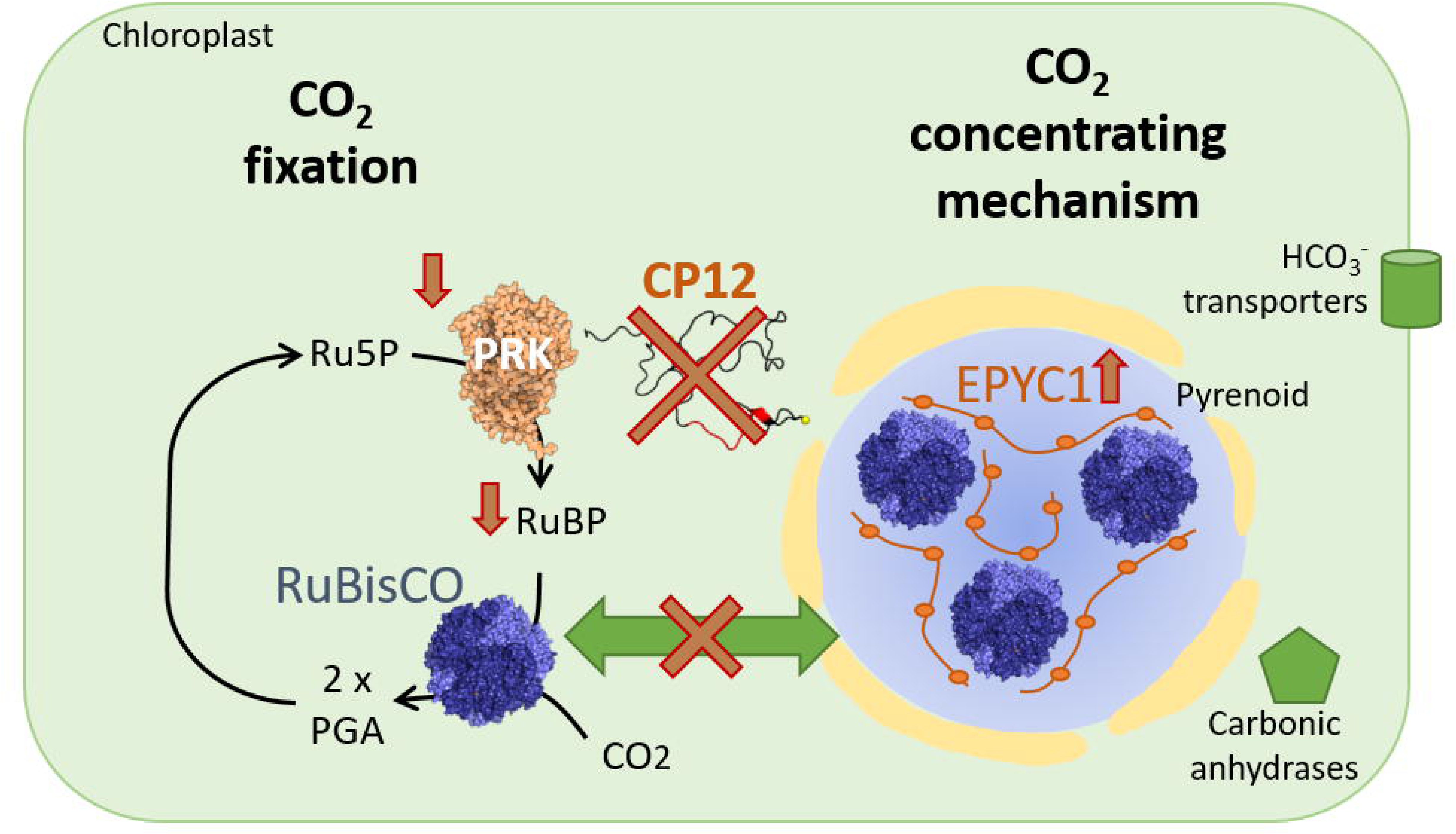

## Introduction

In photosynthetic organisms, CO_2_ fixation through the Calvin-Benson-Bassham (CBB) cycle, also known as the reductive pentose phosphate pathway, involves the carboxylation of ribulose-1,5-bisphosphate (RuBP) by Ribulose 1,5-bisphosphate carboxylase-oxygenase (RuBisCO) (Bassham *et al*., 1950; Andersson and Backlund, 2008). RuBisCO can carboxylate or oxygenate RuBP and the oxygenation reaction leads to photorespiration that reduces productivity (Bowes and Ogren, 1972; Tcherkez *et al*., 2006). The slow diffusion of gases in water can lead to a low ratio of CO_2_ to O_2_ at the active site of RuBisCO, reducing carboxylation in aquatic phototrophs. To circumvent this, many aquatic photosynthetic organisms have evolved biophysical CO_2_ concentrating mechanisms (CCMs) (Kaplan and Reinhold, 1999; Giordano *et al*., 2005; Moroney and Ynalvez, 2007; Long *et al*., 2024). These involve bicarbonate transporters (Duanmu *et al*., 2009a), carbonic anhydrases (Duanmu *et al*., 2009b; Kasili *et al*., 2023) and condensation of RuBisCO in a specific cellular compartment: the pyrenoid in microalgae (Mackinder *et al*., 2016; Fei *et al*., 2022) and the carboxysome in cyanobacteria (Rae *et al*., 2013; Meyer, 2022). The pyrenoid is a membrane-less organelle found in the green microalga *Chlamydomonas reinhardtii* where RuBisCO is condensed through its interaction with the disordered Essential Pyrenoid Component 1 (EPYC1) (Mackinder *et al*., 2016; He *et al*., 2020). The proportion of RuBisCO in the pyrenoid increases at low concentration of CO_2_ (Borkhsenious *et al*., 1998; He *et al*., 2020) or higher concentration of O_2_ (Neofotis *et al*., 2021).

The last step of RuBP regeneration, the phosphorylation of ribulose 5-phosphate by ATP, is catalysed by phosphoribulokinase (PRK) that is regulated by the thioredoxin regulatory network and the small regulatory redox-sensitive protein CP12 (Wedel *et al*., 1997; Oesterhelt *et al*., 2007; Marri *et al*., 2009; Avilan *et al*., 2012; Gontero and Maberly, 2012; Meloni *et al*., 2023). CP12 is a conditionally disordered protein (Graciet *et al*., 2003; Launay *et al*., 2021), and a well-known regulator of the CBB cycle (Gontero and Maberly, 2012). It forms a complex with glyceraldehyde-3-phosphate dehydrogenase (GAPDH) and PRK in the dark reducing their activities (Wedel *et al*., 1997; Lebreton *et al*., 2003; Erales *et al*., 2011; Howard *et al*., 2011b; Avilan *et al*., 2012). Mutants lacking CP12 have smaller amounts of PRK protein in *Arabidopsis thaliana* (López-Calcagno *et al*., 2017) under light-dark cycles and in *C. reinhardtii* under continuous light where the ternary complex is not formed (Gérard *et al*., 2022b; Teh *et al*., 2023). This suggests that CP12 performs functions other than dark inhibition of CBB enzymes (Gontero and Maberly, 2012; Gérard *et al*., 2022a). In addition, in *C. reinhardtii*, the deletion of CP12 also led to a higher amount of EPYC1 compared to wild type cells, even when grown at high CO_2_ (Gérard *et al*., 2022b).

*C. reinhardtii* was chosen as a model organism thanks to the extensive background information available and because it contains only a single copy of the *CP12* gene. *CP12* deleted mutants and partially complemented cells from this alga were used to modulate the abundance of CP12. The abundance of PRK was assessed in these different cell lines, as well as the amount of its product RuBP. The amount of the RuBisCO scaffold protein EPYC1 was also investigated, as well as the RuBisCO location within the pyrenoid or in the chloroplast stroma under high and very low CO_2_ using electron microscopy and immunogold labelling. The acclimation from high to very low CO_2_ in the presence or absence of CP12 was probed using proteomics and by measuring the net photosynthetic activity as function of the dissolved inorganic carbon (DIC) concentration.

## Material and Methods

### Algal strains and their maintenance

*C. reinhardtii* strain CC-4349 (purchased from the Chlamydomonas Resource Center, University of Minnesota, Minneapolis, MN, USA) and Δ*CP12*, (CC-4349 deleted from the protein CP12) described in (Gérard *et al*., 2022b) were maintained on a plate containing Tris Acetate Phosphate (TAP) medium at pH 7.4 ( 20 mM Tris, 0.4 mM KH_2_PO_4_, 0.6 mM K_2_HPO_4_, Beijerinck salts solution containing 7.5 mM NH_4_Cl, 0.4 mM MgSO_4_ and 0.34 mM CaCl_2_ and Hutner’s trace elements (Hutner *et al*., 1950)) and supplemented with agar (15 g L^-1^) and hygromycin (30 µg µL^-1^) for the mutant strain Δ*CP12* and paromomycin (30 µg µL^-1^) for proAR::*CP12* (described below).

### Generation and characterization of a complemented strain of the Δ*CP12* mutant

To restore *CP12* gene expression in the Δ*CP12* mutant, a transgenic vector for *CP12* overexpression was constructed. The pOpt-mVenus-Paro vector was procured from the Chlamydomonas Resource Center (MN, USA) and utilized as the backbone of the vector for overexpression. The vector contains the paromomycin resistance gene, aphVIII, which functions as a selection marker, and the AR promoter for robust expression. The mVenus gene was excised using *Nde I* and *EcoR I* enzymes. Subsequently, a PCR product of the complete *CP12* gene (Cre08.g380250) comprising two introns was generated using KOD Plus (TOYOBO, Japan), and introduced into the mVenus removal site (Fig. S1A). The primers employed are presented in Table S1. Nuclear random transformation was performed via electroporation in accordance with the previously described procedure (Kim *et al*., 2019), utilizing a linearized plasmid that had been digested with the *Pvu II* enzyme. Following transformation, the transformants were selected on TAP agar plates containing paromomycin (10 µg µL^-1^). The suspension of transformed cells was used as a template for PCR with primers specific to proAR::*CP12* (see Table S1) using KOD One master mix (TOYOBO, Japan).

RNA extraction and quantitative RT-PCR methods followed the previously described procedure (Seo *et al*., 2015). Total RNA was extracted using the Hybrid-R RNA purification kit (GeneAll, Korea). cDNA was prepared using the Reverse Transcription Master Premix (ELPIS, Korea). The data are presented as the mean of two technical replicates for each independently prepared biological sample (n = 3), with standard deviation (SD). *CP12* transcript levels were normalized to the expression of the gene encoding RACK1.

### Culture conditions

Pre-cultures were grown in TAP medium supplemented with the appropriate antibiotic for two days under continuous illumination of 110 µmol photon m^2^ s^-1^, photosynthetically available radiation, at 23°C, constantly shaken at 110 rpm.

Then, cells were grown photoautotrophically in medium at pH 7.4 (20 mM MOPS, 0.4 mM KH_2_PO_4_, 0.6 mM K_2_HPO_4_, Beijerinck salts solution containing 7.5 mM NH_4_Cl, 0.4 mM MgSO_4_ and 0.34 mM CaCl_2_ and Hutner’s trace elements (Hutner *et al*., 1950)), illuminated continuously with white LEDs at 110 µmoles of photons m^2^ s^-1^ and at 23°C. Cultures (200 mL in 1 L bottles) were grown under high CO_2_ (HC) by bubbling them with air enriched with CO_2_ at P_CO2_ = 20 000 ppm of CO_2_ resulting in [CO_2_] = 720 µM using a El-Flow Bronkhorst flowmeter with a bubbling rate of 320 mL min^-1^ per litter of culture. This condition was maintained until the culture reached the exponential phase. The cultures were then transferred for 24 hours to very low CO_2_ conditions (VLC). To achieve this, air was passed through soda lime to absorb CO_2_ before being bubbled through the culture, resulting in [CO_2_] ≈ 1.8 µM.

### Protein extraction

Cells in exponential phase were harvested and centrifuged at 2000 g for 10 min at 4°C, then washed in 10 mM Tris at pH 8, containing a protease inhibitor cocktail (Sigma P2714-1BTL). The cell pellet was re-suspended in 1 mL of the same buffer, followed by sonication at an intensity of 40% and with six cycles of 30 s on ice. Protein concentration in solution was determined by the Bradford method (Bradford, 1976), using bovine serum albumin (BSA) as the standard.

### Electrophoresis and immunoblotting

For electrophoresis under denaturing conditions, the proteins were denatured in a buffer including 0.5% of SDS and 20 mM dithiothreitol (DTT) and 30 µg were loaded onto a 12% SDS polyacrylamide gel. Migration was performed at 120 V using a Bio-Rad Mini-PROTEAN system. For native gel electrophoresis, soluble proteins were not subjected to any prior treatment, except the addition of DTT when stated, and electrophoresis were performed using the Phast System with ready-to-use 4–15% acrylamide gradient gels (GE Healthcare). Proteins were then transferred to a nitrocellulose membrane (Trans-Blot Turbo Transfer pack) using a Bio-Rad Trans-Blot Turbo system. After one hour of blocking (5% fat free milk in PBS-Tween20 0.1% buffer), the membrane was incubated overnight with primary antibodies against EPYC1 (Agrisera, diluted 1:5000) or against PRK (antibodies kindly provided by Dr Pierre Crozet (Boisset *et al*., 2023), diluted 1:10000). After washing with PBS-Tween buffer, the membrane was incubated for one hour with a horseradish peroxidase (HRP) coupled anti-rabbit secondary antibodies (Invitrogen, diluted 1:10000). After washing, the HRP substrate (WesternPierce™ ECL Transfer Substrate) was added, and chemiluminescence was detected using a ImageQuant LAS 4000 mini.

### Metabolite extraction

The extraction protocol is the same as described in (Saint-Sorny *et al*., 2022), with a few modifications. 25 mL of exponentially growing *C. reinhardtii* culture at HC (see culture conditions) were quenched in 50 mL of cold 70% methanol. The number of cells per mL was estimated using the LUNA-II™ automatic cell counter. All steps of the extraction process were done in an ethylene glycol/ethanol solution (80/20) bath at −20°C. 5 mL of chloroform was then added to 25 mL of the quenched culture. A Potter glass homogeniser was used to homogenize the mixture, followed by three freeze/thaw cycles to ensure complete cell lysis. The extracts were centrifuged at 2851 g at 4°C for 10 min to collect the aqueous phases containing metabolites. The organic phase was washed twice with ice-cold water, and these washing fractions were combined with the aqueous phase to be lyophilized.

The freeze-dried fractions were resolubilized in 1.5 mL H_2_O. The resulting supernatant was cleaned up with two solid-phase extractions, using the columns Strata-C and Strata-X-AW (Phenomenex, Le Pecq, France). 10 μL H_3_PO_4_ 50% were added to the extract that was then loaded on the Strata-X-C column (flow rate 1 mL min^-1^) after conditioning with methanol and water. The loading fraction was collected in a tube. The column was washed with 1 mL H_3_PO_4_ 0.1% and the washing fraction was collected. Elution was carried out with NH_3_/MeOH/H_2_O (0.5 mL 5/25/70 v/v/v, 0.5 mL 10/25/65, 1 mL 20/25/55; flow rate 1 mL min^-1^). The eluate was collected in an Eppendorf tube. Both loading and washing fractions were gathered and 500 mL phosphate buffer (0.01 M) was added. The resulting buffered fraction was loaded (1 mL min^-1^) on the Strata-X-AW column after conditioning with pure methanol, methanol/formic acid/water (2/25/73 v/v/v) and then phosphate buffer. The column was then washed with 1 mL phosphate buffer. Elution was carried out with NH_3_/MeOH/H_2_O (0.5 mL 5/25/70 v/v/v, 0.5 mL 10/25/65, 1 mL 20/ 25/55; flow rate 1 mL min^-1^). The eluate was collected in an Eppendorf tube. The two eluates (C and AW) were spin-dried (speed-vac) and used immediately for LC-TOF analysis.

### RuBP quantification

For LC-TOF analyses (Guérard *et al*., 2011), the spin-dried extracts were resuspended in 100 μL milli-Q water, centrifuged (8000 g, 4°C, 20 min) and the supernatant was transferred to a glass vial. Samples were injected in the UHPLC 1290 infinity II (Agilent Technologies, Les Ulis, France) equipped with the column UPLC-HSS T3 (2.1 × 100 mm, 1.8 mm) (Waters, Guyancourt, France) under an ammonium acetate/methanol gradient (99/1-0/100%, 6 min). The UHPLC was connected to the injection orifice of the mass spectrometer via a capillary tube. The flow rate was 0.4 mL min^-1^.

Electrospray (Jet Stream Technology Ion Source) mass spectrometry was carried out with the 6540 Q-TOF (Agilent Technologies, Les Ulis, France) with N_2_ as a spray gas, at 325 °C (Gas Temp), 10 L.min^-1^ (Drying Gas), 33 psi (Nebulizer), 400 °C (Sheath Gas Temp) and 12 L min^-1^ (Sheath Gas Flow), upon negative source ionization (3500 V capillary voltage). Resolution parameters were adjusted to achieve optimal mass resolution in the mass range 50-1100 amu. Mass spectra were recorded with an acquisition rate of 2 spectra s^-1^ and acquisition time 500 ms per spectrum. Data were acquired and processed using the software MassHunter Data Acquisition, MassHunter Qualitative Analysis and MassHunter Quantitative Analysis. All compounds analysed here were measured and calibrated with pure standards purchased at Sigma-Aldrich/Fluka (Saint-Quentin Fallavier, France).

### Immunolabelling Transmission Electron Microscopy and image analysis

*C. reinhardtii* cells cultured at HC and VLC (see culture conditions) were pelleted, high pressure frozen, freeze substituted and embedded in LR White resin (Hard Grade), according to (McDonald, 2014). Ultrathin sections were prepared with an ultracryomicrotome (EM UC7 Leica), and immunogold labelling was performed (O’Toole, 2010). Primary antibodies RuBisCO raised against the large subunit (antibodies kindly provided by Dr Xenie Johnson, dilution 1/500) were used to detect RuBisCO, and were coupled to secondary Aurion Protein G conjugated to 10 nm gold particles (nanogold particles). The samples were analysed using a Tecnai 200 kV electron microscope (Thermofisher Scientific), and digital acquisitions were made with a numeric camera (Oneview, Gatan).

The ‘Advanced Weka Segmentation’ plugin from Fiji (ImageJ 1.54f, available in supplementary data 1) was used to train a model to detect gold particles in cell images. Images containing cross section through a pyrenoid were selected. A mask was generated for each strain and treatment. A Fiji macro was developed to define the pyrenoid region and the entire cell area (in µm^2^). Two regions of the cell were considered for the analysis: the pyrenoid and the entire cell minus the pyrenoid. The macro reported the number of particles counted and the area of these two compartments. The proportion of particles in the pyrenoid was calculated from the number of counts in the pyrenoid divided by the number of counts in the entire cell. The density was calculated from the number of counts divided by the area.

### Proteomic analysis

#### Samples preparation for LC-MS/MS analysis

Extracted proteins (30 µg) were loaded onto a 12% SDS polyacrylamide gel (Bio-Rad Mini-PROTEAN system) and migration was stopped just after the samples entered the separation gel. After staining the gel with Coomassie Blue R250, the bands containing the total proteins were cut and washed twice in a mixture of 100 mM NH_4_HCO_3_ and CH_3_CN (50/50, v/v) for 10 min at 37°C with 400 rpm agitation. The samples were then reduced with 10 mM DTT for 45 min at 56°C in the dark, followed by alkylation with a solution of 55 mM iodoacetamide for 30 min at room temperature in the dark. The bands were washed again as described above. Then, 80 μL of CH_3_CN solution was added, and after its evaporation, the samples were digested with trypsin/LysC protease (10 to 20 ng in 25 mM NH_4_HCO_3_) overnight at 37°C. The tryptic peptides were extracted twice with 0.1% TFA in water/ CH_3_CN (50/50, v/v). The supernatants containing tryptic peptides were recovered by centrifugation, pooled and dried for 2 h, at 45°C. About 1 µg of digested peptides from each condition was injected on a nanoliquid chromatography coupled to Q-Exactive plus mass spectrometer (Thermo Electron) as previously described in (Gérard *et al*., 2022b) except for the nanochromatography which was a NeoVanquish system from Thermofisher and the separation of peptides was performed on a EASY-Spray PepMap Neo C18 (1500 bar, 75 µm x 150 mm) using a similar gradient. The same top 10 Data dependent acquisition mode was applied.

### Identification and label free relative quantification

Data from six biological replicates were processed by the computational proteomics platform MaxQuant (version 2.4.14) for protein identification and quantification as previously described in (Gérard *et al*., 2022b) with the database of *C. reinhardtii* CC-4532 v6.1 (NCBI-Taxonomy-ID = 3055) extracted from Phytozome v13 (30 678 proteins). Perseus software (version 1.6.15.0) was used for statistical analysis. The data were filtered to eliminate contaminants, false positives, proteins that are only identified by peptides that carry one or more modified amino acids. LFQ (label-free quantification) intensities were transformed to logarithm base 2 (log2) to follow a normal distribution. Quantifiable proteins (valid values) were defined as those detected in at least 70% of samples within at least one group, resulting in a total of 2641 proteins for analysis. Missing values were replaced by low values following a normal distribution. A t-test with a False Discovery Rate threshold of 0.05 and an S0 value of 0.1 was performed to identify proteins that differed significantly between two groups and volcano plot representation was designed using R Studio (Version 2024 12.1). To determine the exact pairwise differences of protein expression Tukey’s Honest Significant Difference (THSD) test was used on the ANOVA significant hits (with a False Discover Rate threshold of 0.05) in Perseus. Heat map representation was designed using R Studio (Version 2024 12.1).

### CP12 targeted proteomics

For targeted quantification of CP12, a list of CP12 tryptic peptide m/z was included in the MS/MS data acquisition program (Table S2), to focus on CP12. Data were processed on Proteome Discoverer 3.0 (Thermofisher, SequestHT algorithm), with a static modification for carbamidomethylation on Cys (+57.021 Da). Quantification of CP12 was performed by spectral counting based on Peptide Spectral Matches.

### EPYC1 phosphorylation identification

For the identification of the phosphorylated amino acid residues of EPYC1, data from three biological replicates were processed on Proteome Discoverer 3.0. A specific processing workflow for phosphorylated peptides was used including the SequestHT and MS Amanda algorithms with Percolator validation and the IMP-ptm-RS node for phosphorylation site probability. Dynamic modifications were searched for oxidation on Met (+15.995 Da) and phosphorylation on Ser/Thr/Tyr (+79.966 Da).

### Photosynthetic activity

Cells from three independent cultures at HC or VLC (see culture conditions) were centrifuged at 2000 g for 5 min and washed twice with the culture MOPS medium described above at pH 6.9 or 7.9 and degassed by flushing with N_2_. These pH values are each only 0.5 units from the growth pH but produce different proportions of CO_2_ to HCO_3_^-^: the ratio of CO_2_ to HCO_3_^-^ is 0.25 at pH 6.9, but 0.025 at pH 7.9 (Dickson and Millero, 1987).

The net rate of photosynthesis was measured as the evolution of oxygen (Oxygraph; Hansatech Instruments, UK) using the Oxygen View software. Cells (2 mL at 10^6^ cells mL^-1^) were placed in the Oxygraph chamber and illuminated with 110 µmol photon m^2^ s^-1^, at 23° C. When the steady state rate reached a plateau, increasing amounts of sodium bicarbonate (NaHCO_3_) were added to produce a range of DIC, from 0.012 to 2 mM. The rate of O_2_ released was measured for each concentration and the data, triplicated for each condition, were fitted to a pseudo Michaelis-Menten equation using SigmaPlot v14 (equation 1).

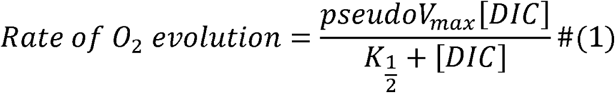

Differences in the proportions of CO_2_ and HCO_3_ ^-^ at the two pH values were used to discriminate between uptake of these two forms using a model (equation 2 and Clement *et al*., 2016) where α represents the contribution of CO_2_ to the total rate of O_2_ evolution.

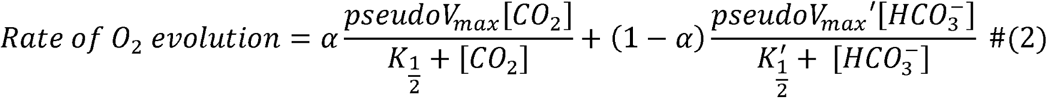

The rates were normalised by the amount of chlorophyll *a* as previously described (Clement *et al*., 2016). The chlorophyll *a* total quantity in the Oxygraph reaction chamber ranged from 9 to 12 µg. To quantify the chlorophyll *a*, cells were centrifuged at 2000 g for 5 min and the cell pellet was resuspended in 90% of acetone, incubated for 30 min at 4°C in the dark and centrifuged for 10 min at 10,000 g at 4°C. Absorbance of the supernatant was measured at 630 nm, 647 nm and 664 nm and chlorophyll *a* quantified using the equations of (Ritchie, 2008).

### Statistical Analysis

Data analysis and visualization were performed using R with theggplot2 package (Wickham, 2016; R Core Team, 2021). The Kruskal-Wallis test, followed by Dunn’s post hoc test was used for multiple comparisons for microscopy data (Kruskal and Wallis, 1952; Dunn, 1964).

## Results

### CP12 abundance in different cell lines

The abundance of CP12 in different cell lines was probed by a targeted proteomic approach. No CP12 peptides were detected in the Δ*CP12* deletion mutants (Fig. 1A). A partially complemented Δ*CP12 [proA::CP12]* mutant (named Δ*CP12::CP12* hereafter) was generated from the strain Δ*CP12* by reintroducing the *CP12* gene (Fig. S1). The CP12 level in Δ*CP12::CP12* was half that of wild type (WT) cells (Fig. 1A). As expected regarding the known function of CP12, the ternary complex of GAPDH, PRK and CP12 was not observed in Δ*CP12* under oxidizing conditions but was present in Δ*CP12::CP12* (Fig. S2) even though the abundance of CP12 in the Δ*CP12::CP12* was lower than in the WT. The abundance of CP12 in WT cells did not change with the CO_2_ concentration of the cell culture (Fig. S3A).

**Fig. 1.**
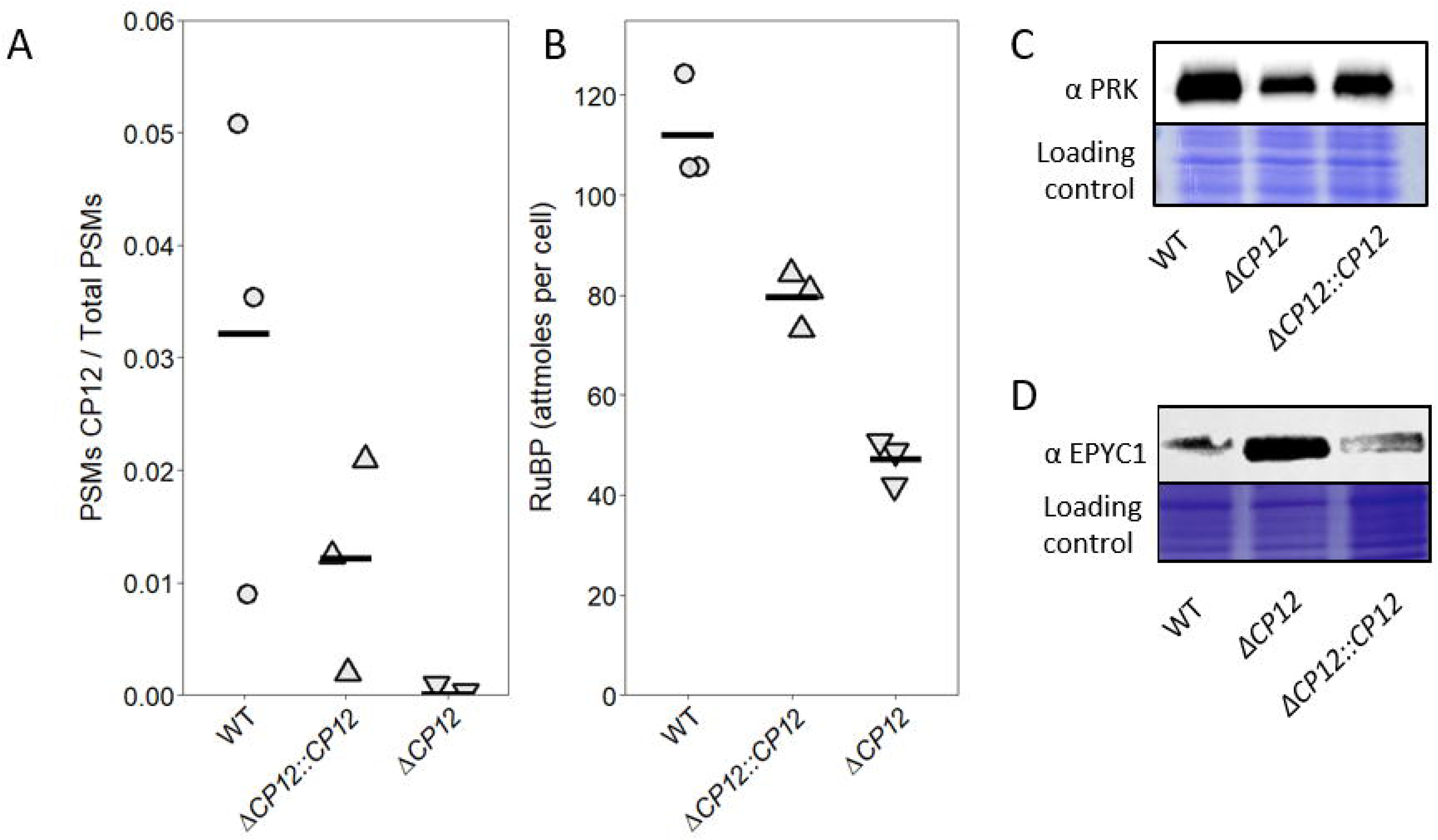
Abundance of PRK, EPYC1 and RuBP in *C. reinhardtii* strains with varying CP12 levels. (A) Peptide Spectrum Matches (PSMs) corresponding to CP12 in WT, Δ*CP12::CP12* and Δ*CP12* grown at high CO_2_ (HC). PSMs of CP12 were obtained using targeted mass spectrometry and normalized by the total PSM count. (B) RuBP quantification (attomoles cell^-1^) by LC-MS/MS in WT, Δ*CP12::CP12* and Δ*CP12* grown at high CO_2_ (HC). In (A) and (B), symbols show individual replicates and the horizontal lines show their mean (n=3, technical replicates). (C) (D) Western blot analysis of PRK (C) and EPYC (D) in total protein extracts from WT, Δ*CP12* and Δ*CP12::CP12* strains grown at HC. Coomassie blue stain of the SDS-PAGE was used as loading control.

### Effect of CP12 level on the amount of PRK and RuBP

Western blots were used to determine the relative amount of PRK in the strains with different levels of CP12. In cells grown at HC, the amount of PRK was lowest in Δ*CP12,* increased in Δ*CP12::CP12*, and was highest in WT cells (Fig. 1C). The amount of the PRK product, RuBP, was 2.4-fold lower in Δ*CP12* than in the WT (Fig. 1B). In Δ*CP12::CP12*, the amount of RuBP was 1.4-fold lower than in WT cells, similar to the trend for CP12 and PRK levels. The relative amount of PRK was also probed as a function of CO_2_ in three independent mutants Δ*CP12* (Fig. S3B). Its level was lower in Δ*CP12* than in WT at both CO_2_ concentrations. The abundance of PRK decreased from HC to VLC in WT strain.

### Effect of CP12 level on the amount of EPYC1 and its phosphorylation state

Western blots were used to determine the relative amount of EPYC1 in the strains with different levels of CP12 (Fig 1D). The amount of EPYC1 was higher in Δ*CP12* than in WT in cells grown at HC (Fig. 1D). In Δ*CP12::CP12*, the amount of EPYC1 was similar to that in WT cells at HC. The abundance of EPYC1 was also probed as a function of CO_2_ in WT and in three independent mutants Δ*CP12* (Fig. S3A). The amount of EPYC1 was higher in VLC than in HC in WT and Δ*CP12* mutants, and was higher in Δ*CP12* mutants than in WT at both CO_2_ concentrations (Fig. S3B). Since EPYC1 is phosphorylated (Turkina *et al*., 2006), its phosphorylation state was also analysed in WT and Δ*CP12* strains grown at HC and upon acclimation to VLC for 24 h. Several serine and threonine residues within repeat motifs of EPYC1 were phosphorylated regardless of the CO_2_ growth conditions in the WT and the three independent Δ*CP12* mutants (Fig. S4), such as Ser137 within the second RuBisCO Binding Site (RBS), Ser197 within the third RBS and Ser258 within the fourth RBS (numbering from the pre-protein including the chloroplast targeting peptide). The residues Ser53 of the first RBS, Thr116 of the second RBS, Thr176 of the third RBS and Thr237 of the fourth RBS were found to be phosphorylated in the WT at HC and VLC as well as in the three independent Δ*CP12* at HC. However, the peptides containing these residues were not found in the proteomic analysis of the Δ*CP12* at VLC.

### Effect of *CP12* deletion on RuBisCO location

EPYC1 promotes RuBisCO location to the pyrenoid, therefore the distribution of RuBisCO was examined in WT, Δ*CP12*, and Δ*CP12::CP12* lines to assess its relationship with EPYC1 and CP12 levels. Transmission electron microscopy images showed that the cellular ultrastructure was similar in all strains and treatments (Fig. 2A and Fig. S5). In contrast, the proportion of RuBisCO assessed by immunogold-labelling inside and outside the pyrenoid for each strain was different (Fig. 2B, C, D). In the WT, the proportion of RuBisCO outside the pyrenoid decreased from 0.64 at HC to 0.22 at VLC (Fig. 2B, Table S3). Δ*CP12::CP12* behaved similarly with a change of proportion from 0.51 to 0.26. In contrast, in Δ*CP12*, the difference of proportion of RuBisCO inside and outside the pyrenoid was less pronounced at HC and VLC (0.37 to 0.46, Fig. 2B).

**Fig. 2.**
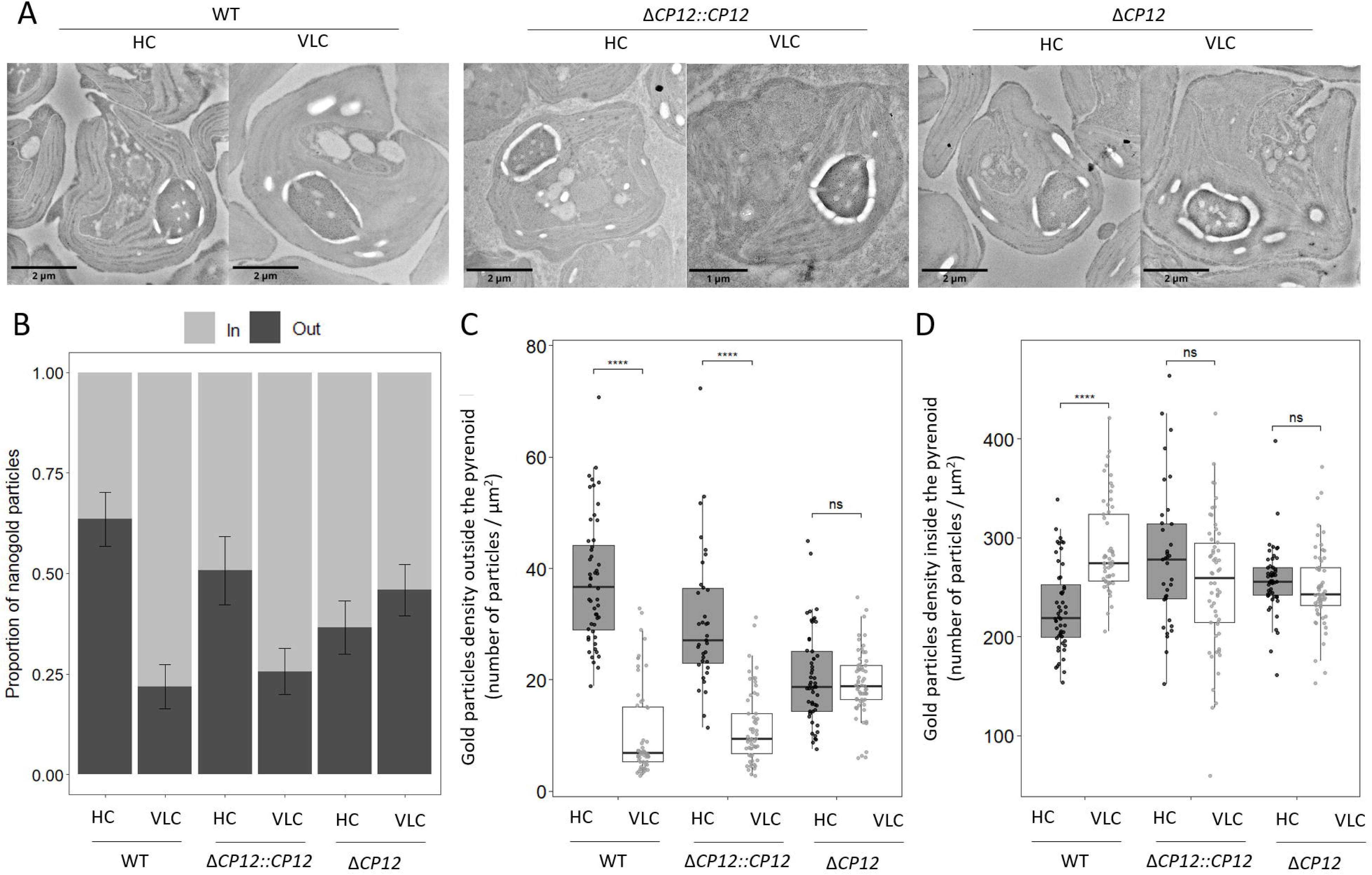
Cellular ultrastructure and location of RuBisCO probed by immunogold labelling. (A) Representative transmission electron microscopy images of cells grown at HC and after 24 hours at VLC for the three strains; more images are shown in Fig. S5. (B) Proportion of gold particles inside and outside the pyrenoid. (C and D) Density of gold particles (number of particles per µm^2^) outside (C) and inside (D) the pyrenoid. The statistical significance was computed by the Kruskal-Wallis test, followed by Dunn’s post hoc test for multiple comparison (ns, p > 0.05 and ****, p < 0.0001). The lower and upper hinges correspond to the first and third quartiles. The upper and lower whiskers extend from the hinge to 1.5 times the inter-quartile range (n averages 53 and ranges from 35 to 61, Table S3).

To account for possible bias introduced by the position of the sections through the pyrenoid, the data were interpreted in terms of areal density (Fig. 2C and D). In WT cells, the density of RuBisCO outside of the pyrenoid dropped from 37.7 to 10.4 particles µm^-2^ upon transition from HC to VLC. A similar decrease was observed for Δ*CP12::CP12* (from 30.5 to 11.4 particles µm⁻²). In contrast, the density outside the pyrenoid remained unchanged for Δ*CP12* under both HC and VLC with a density around 20 particles µm^-2^. The density of RuBisCO inside the pyrenoid was less variable than outside among the three strains and CO_2_ treatments, and only in the WT cells was RuBisCO more densely packed at VLC than at HC.

### Differential protein abundance in WT and CP12 deficient strains upon transition from HC to VLC

Well-known CCM candidates and low CO_2_ inducible (LCI) proteins (Grossman and Wollman, 2023) were quantified using label-free relative quantitative proteomics. Changes in protein abundance were probed for both WT and Δ*CP12* strain upon a transition from HC to VLC (Fig. 3A and B, Fig. S6 and Supplementary data 2). In both strains, CCM components and LCI proteins were the most differentially abundant in VLC compared to HC. In the WT, 398 proteins were differentially abundant, with 163 being more abundant in VLC compared to HC. In the mutant, 112 proteins were differentially abundant, including 56 more abundant at VLC (Supplementary data 3). 77 proteins were common in the two sets of 398 and 112 proteins (Fig. 3C). Among these, 37 proteins were more abundant at VLC in both WT and Δ*CP12* strains (Supplementary data 3). These included HCO_3_ ^-^ transporters known to be involved in CCM: LCI1 (Cre03.g162800) localised at the plasma membrane (Kono and Spalding, 2020), CCP1 (Cre04.g223300) and NAR1B (Cre06.g309000, also named LCIA) both localised at the chloroplast membrane (Pollock *et al*., 2004; Wang and Spalding, 2014; Förster *et al*., 2023). The LCIC1 chloroplastic carbonic anhydrase (Cre06.g307500) and its homologue LCID1 (Cre04.g222800) (Yamano *et al*., 2022) were also among these 37 proteins, as well as a mitochondrial carbonic anhydrase CAH5 (Cre05.g248450) (Rai *et al*., 2021). Other low CO_2_ inducible proteins were also among these 37 proteins, such as: LCI19, a glyoxylate reductase/ 6-phosphogluconate dehydrogenase (Cre11.g467617); LCI9 that contains a starch binding domain (Cre09.g394473); light-harvesting complex I chlorophyll a/b binding protein LHCSR3A (LHCA1, Cre08.g367500) and LCI23 (Cre06.g281600) of unknown function.

**Fig. 3.**
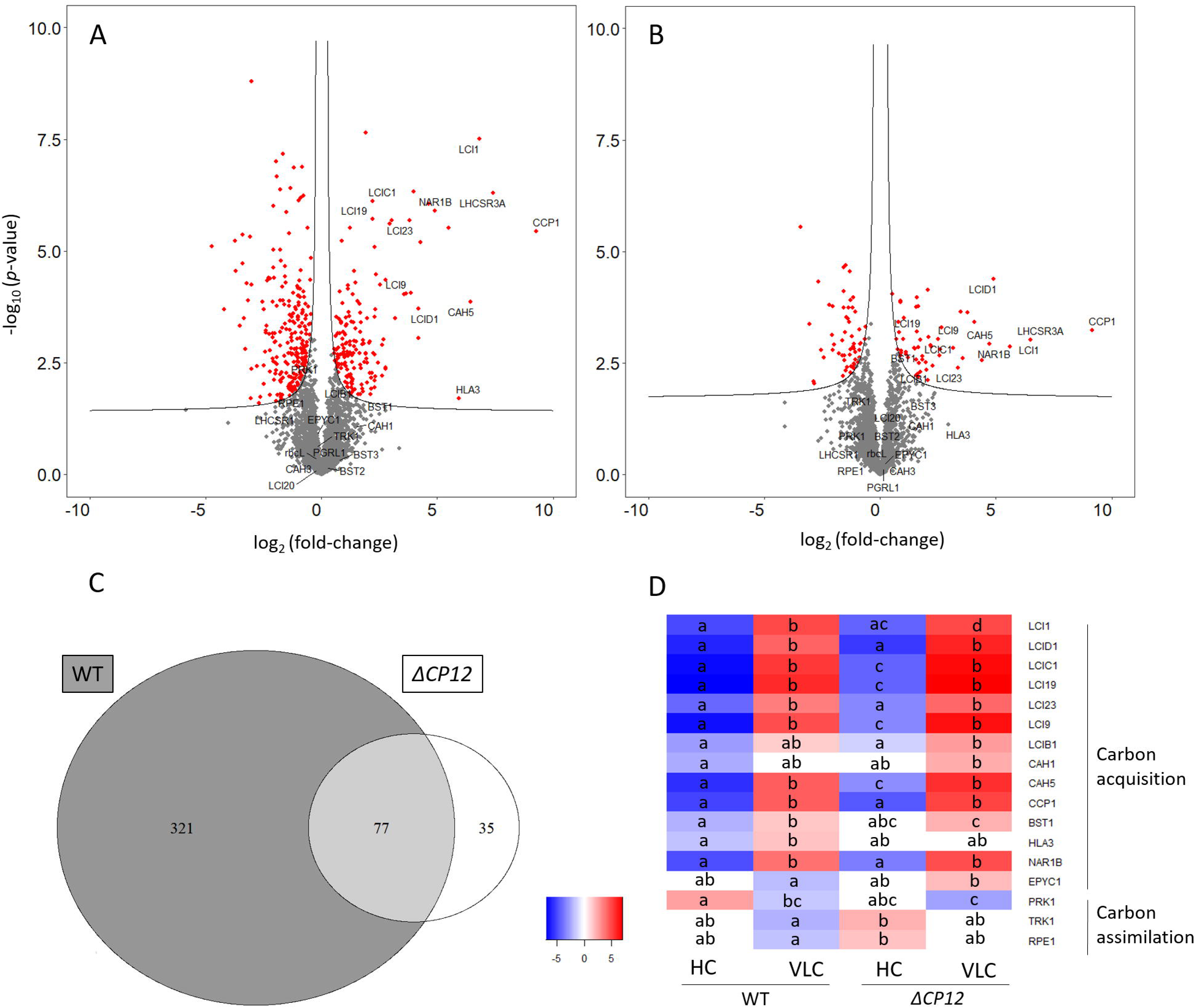
Differential protein abundance in WT and Δ*CP12* strains on acclimation from HC to VLC. (A) and (B): Volcano plots showing the changes of abundance of proteins in WT (A) and Δ*CP12* (B) cells after 24 hours acclimation to VLC conditions compared to HC conditions. The x-axis shows the change in protein abundance on a log_2_ scale 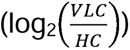 while the y-axis shows the statistical significance (–log₁₀ p-value). The identifiers for all these proteins are included in Table S4. Proteins above the line are significantly different and are shown in red. Other volcano plots comparing WT *vs* Δ*CP12* are represented in Fig. S6. (C): The Venn diagram illustrates the number of proteins that changes in abundance significantly on acclimation from HC to VLC, based on a t-test with a false discovery rate of 0.05. (D): The heatmap represents the abundance of proteins involved in carbon acquisition and assimilation in WT and Δ*CP12* at HC or VLC. Different letters (a, b, c, d) denote significant pairwise differences for each protein, resulting of a post hoc Tukey’s HSD test following multiple ANOVA. The protein abundances (on a log_2_ scale) are represented in heatmaps Fig. S7 (proteins involved in carbon acquisition and assimilation) and Fig. S8 (all proteins).

ANOVA tests followed by post hoc analysis were used to analyse the effects of both CP12 levels and CO_2_ concentration on proteins abundance (Fig. 3D, Fig. S7, S8 and Supplementary data 4 and 5). In this analysis, the bicarbonate transporter of the plasma membrane HLA3 (Cre02.g097800) (Duanmu *et al*., 2009a) responded differently in the two strains: it was more abundant in VLC compared to HC in the WT but did not change in the mutant.

PRK abundance was lower in the WT at VLC compared to HC, as was also found with the Western blot. The higher abundance of EPYC1 in Δ*CP12* compared to WT was only observed at VLC. Other proteins of the CBB cycle were not significantly different among strains and CO_2_ conditions.

### Effect of *CP12* deletion on inorganic carbon acquisition kinetics

The rates of net photosynthesis of *C. reinhardtii* cells grown at HC or after 24 hours of acclimation at VLC were measured as a function of DIC concentration at pH 6.9 and pH 7.9 (Fig. 4 and Table S5). For both WT and Δ*CP12* strains, the K_1/2_ for DIC was almost two-fold lower after 24 hours at VLC compared to HC, at both pH values (Fig. 4, Table S5). However, for both WT and Δ*CP12,* the pseudo-V_max_ was not significantly different between HC and VLC at both pH values.

**Fig. 4.**
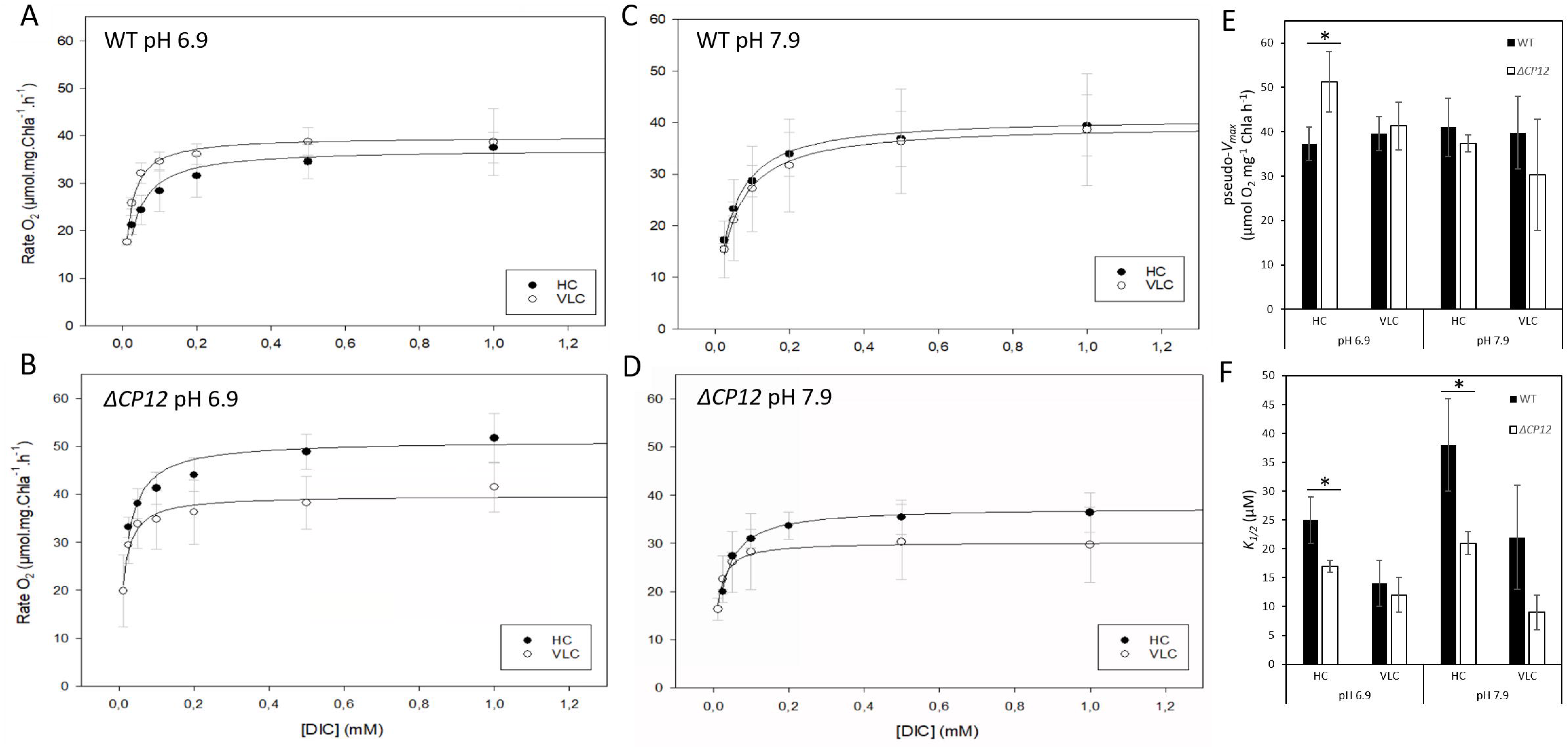
Kinetics of photosynthesis of WT and Δ*CP12* at HC and at VLC. Rate of net photosynthesis for cells grown at HC (black circles) and after 24 hours of acclimation to VLC (white circles) for the WT (A) and (C) and the mutant Δ*CP12* (B) and (D), and measured at pH 6.9 (A) and (B) and at pH 7.9 (C) and (D). (E) Histogram of the fitted pseudo-V_max_ for the different samples. n=3, biological replicates, unpaired t-test (*, p < 0.05) (F). Histogram of the fitted K_1/2_ for the different samples using the Michaelis– Menten equation (equation 1).

Comparing the two strains at VLC, the K_1/2_ and pseudo-V_max_ were not significantly different at both pH values. At HC, in contrast, the K_1/2_ for DIC for Δ*CP12* cells was significantly lower than for the WT when measured at both pH values. The pseudo-V_max_ for Δ*CP12* at HC was significantly higher than the WT at pH 6.9 but not at pH 7.9. To analyse further the difference in response between WT and Δ*CP12* cells grown at HC, kinetic parameters were also modelled as a function of CO_2_ and HCO ^-^ concentration and the K_1/2_ for HCO_3_ ^-^ decreased from 84.0 µM for the WT to 0.1 µM for the Δ*CP12* (Table S6).

## Discussion

### Acclimation of WT strain from HC to VLC

*C. reinhardtii* is the traditional model for CCM and pyrenoid studies and its CCM components have been well described (for literature reviews, see among other (Mackinder, 2018; Grossman and Wollman, 2023)). The CCM can involve several bicarbonate transporters (Pollock *et al*., 2004; Duanmu *et al*., 2009a) and carbonic anhydrases (Van and Spalding, 1999; Duanmu *et al*., 2009b; Wang and Spalding, 2014; Kasili *et al*., 2023) that together mediate the transport of inorganic carbon towards the pyrenoid and concentrate CO_2_ at the active site of RuBisCO. RuBisCO can be located both in the stroma and the pyrenoid, and at VLC, it is mostly present in the pyrenoid of *C. reinhardtii* (Borkhsenious *et al*., 1998). For WT cells, the proportion of pyrenoid-localised RuBisCO at HC was similar in this study to that described in Borkhsenious *et al*., i.e. 40%. Upon a transition to VLC, Borkhsenious *et al*. found 89% of RuBisCO located in the pyrenoid, and in our study, it was 78%. The shift from HC to VLC also resulted in a lower K_1/2_ for DIC in WT cells grown at VLC than at HC. At HC, the K_1/2_ for DIC measured in this study was very close to that found by Borkhsenious *et al*. (25 µM at pH 6.9 and 19 µM at 7.3) but at VLC, it was higher than that measured by Borkhsenious *et al* (14 µM at pH 6.9 vs <5 µM at pH 7.3, respectively). In the literature, K_1/2_ for DIC for *C. reinhardtii* cells decreased by a factor of 5 to 6 upon a transition from HC to VLC (Borkhsenious *et al*., 1998; Yamano *et al*., 2015; Findinier *et al*., 2024) while here the K_1/2_ only varied 2- to 3-fold at pH 6.9 and 7.9 respectively. The higher K_1/2_ and lower proportion of pyrenoid-localised RuBisCO found here for WT at VLC compared to the literature could explain the differences at VLC.

CCM was induced at VLC in our culture conditions since known CCM hallmarks (Wang *et al*., 2015). were in higher abundance compared to their abundance at HC for WT. Previously described hallmarks we found in our study include chloroplast carbonic anhydrase LCIC1 (Yamano *et al*., 2022), mitochondrial CAH5 (Rai *et al*., 2021), the bicarbonate transporters of the plasma membrane LCI1 and HLA3 (Duanmu *et al*., 2009a; Kono and Spalding, 2020), and of the chloroplast membrane CCP1 (Pollock *et al*., 2004) and NAR1B (or LCIA) (Wang and Spalding, 2014; Förster *et al*., 2023) and of the thylakoid membrane BST1 (Cre16.g662600) (Mukherjee *et al*., 2019). Of interest the abundance of the periplasmic chloroplast carbonic anhydrase LCIB1 (Cre10.g452800), that forms a complex with LCIC1 (Yamano *et al*., 2022), did not change with CO_2_ conditions as much as its partner. The abundance of the periplasmic carbonic anhydrase CAH1 (Cre04.g223100) (Shimamura *et al*., 2024), was also less sensitive to CO_2_ conditions compared to the other CCM hallmarks mentioned above. A range of low inducible proteins were also more abundant at VLC compared to their abundance at HC, indicating the acclimation to VLC in the WT strain.

Our data indicate that, in the WT strain, the shift in proportion of pyrenoid-localised RuBisCO from HC to VLC was largely the result of a lower RuBisCO density in the stroma at VLC, whereas there was limited variation in density within heavily dense RuBisCO biocondensates. The condensation of RuBisCO is described as the result of transient and multiple interactions between EPYC1 and RuBisCO (Freeman Rosenzweig *et al*., 2017; Wunder *et al*., 2018), resulting in very densely packed RuBisCO particles as it was observed here.

### Acclimation of Δ*CP12* strain from HC to VLC

On adaptation to HC or VLC, the amount of CP12 protein did not change significantly. This was consistent with transcriptomic data upon transition from HC to VLC (Fang *et al*., 2012). As previously shown, at HC, the absence of CP12 induced an increase of EPYC1 compared to the WT strain (Gérard *et al*., 2022b). Here, we show that in Δ*CP12*, RuBisCO density in the stroma was lower than that of WT at HC, and higher than the WT at VLC. The higher the level of EPYC1, the higher was the proportion of RuBisCO in the pyrenoid, consistent with EPYC1 being a “molecular glue” for RuBisCO (Wunder *et al*., 2018; He *et al*., 2020). Two studies have identified four RBS in EPYC1 (He *et al*., 2020; Meyer *et al*., 2020) (Fig. S4), and we found that the phosphorylation pattern of these EPYC1 RBS was identical between WT and Δ*CP12* at HC. Turkina *et al*. found that the 2^nd^ and 3^rd^ RBS were phosphorylated at LC but not at HC (Turkina *et al*., 2006). The difference between their data and ours could result from variation in culture conditions as discussed by Turkina *et al.,* as many factors can affect phosphorylation patterns besides CO_2_ conditions. Further understanding of the relationships between the phosphorylation state of EPYC1 and its ability to condense RuBisCO in the pyrenoid is required to rationalise this difference.

Besides, without CP12, the density of RuBisCO in the stroma did not change with CO_2_ availability, as if the signal for RuBisCO relocation was missing. The absence of relocation of RuBisCO towards the pyrenoid at VLC is a unique feature of the Δ*CP12* strain, and it is puzzling that this does not prevent CCM from being activated in Δ*CP12* as seen by lower K_1/2_ for DIC at VLC compared to that at HC and by the induction of most CCM hallmarks mentioned above.

Interestingly, the total number of proteins that changed from HC to VLC was significantly lower in Δ*CP12* compared to WT suggesting that the change in CO_2_ affected the Δ*CP12* strain less than WT strain. Several transcription factors have been identified to be involved in CCM activation such as CCM1/CIA5 or LCR1 (Fukuzawa *et al*., 2001; Yoshioka *et al*., 2004). Since the proteins encoded by the target genes of these transcription factors were present, it seems unlikely that CCM1/CIA5 and LCR1 are impaired. Interestingly, the difference in abundance of HLA3 as a function of CO_2_ was lower in the Δ*CP12* strain compared to the WT strain. *HLA3* has been reported to be regulated by the bZIP transcription factor BLZ8 (Choi *et al*., 2021) and other complex regulatory pathways (Yamano *et al*., 2015; Wang *et al*., 2016), that could be affected by the CP12 deletion. However, HLA3 is proposed to work cooperatively with NAR1B (Duanmu *et al*., 2009a; Wang and Spalding, 2014; Wang *et al*., 2015; Yamano *et al*., 2015), which change in abundance was similar in Δ*CP12* and WT in contrast to its partner.

Altogether, RuBisCO relocation to the stroma under HC conditions or to the pyrenoid under VLC conditions was prevented and a weaker remodelling of the proteome at VLC was induced in the mutant compared to the WT. Despite this, the CCM was still active in Δ*CP12* at VLC. The Δ*CP12* strain is a unique example where RuBisCO location is not associated with the induction of a CCM.

### Differential CO_2_ acquisition mechanisms between Δ*CP12* and WT

At HC in the Δ*CP12* strain, the K_1/2_ for DIC was lower that of WT. The Δ*CP12* strain was able to use HCO_3_^-^ and CO_2_ at HC and had a lower K_1/2_ for HCO_3_^-^ compared to CO_2_, while the WT had a lower K_1/2_ for CO_2_ compared to HCO_3_^-^. The lower K_1/2_ for HCO_3_^-^ in Δ*CP12* than in WT is possibly a consequence of a greater amount of EPYC1 and RuBisCO located in the pyrenoid, rather than being due to an increase in the amount of bicarbonate transporters and carbonic anhydrases at HC. Indeed, there was no relationship between RuBisCO condensation in the pyrenoid and the induction of other CCM components as discussed above. Although this has not been reported previously, it is in line with the theoretical modelling of contribution of different CCM components in *C. reinhardtii* (Fei *et al*., 2022).

The question remains, however, how inorganic carbon is transported to the pyrenoid, and how this is energised when CCM components are not induced at HC. The complete re-routing of the *C. reinhardtii* metabolism observed previously in the Δ*CP12* at HC (Gérard *et al*., 2022b), including metabolic pathways connected by the malate valves (Dao *et al*., 2022), could contribute to this adaptation. Deletion of *CP12* in tobacco resulted in lower malate valves capacity (Howard *et al*., 2011a). Among other proteins, the abundance of the mitochondrial carbonic anhydrase CAH5 was higher in the Δ*CP12* than in the WT at HC. CAH5 has been suggested as an important cross-compartment coordination between mitochondria and chloroplasts (Burlacot and Peltier, 2023; Wang *et al*., 2023; Findinier *et al*., 2024), and could contribute to the rerouting of the metabolic flux. Furthermore, the mitochondrial relocation that was typically observed upon a transition from HC to VLC was absent in a deficient *PRK* mutant strain (Boisset *et al*., 2023; Findinier *et al*., 2024).

This difference in DIC acquisition between the two strains was observed at HC. At VLC, the CCM was active in both strains, even though there was a large difference in RuBisCO location, as discussed above.

### Effect of CP12 on RuBP regeneration

The lower levels of CP12 led to a decrease in PRK and RuBP levels but this did not alter the growth of *C. reinhardtii* regardless of the CO_2_ concentration. This is in line with previous studies that indicated that a reduction of PRK amount or activity is not detrimental to photosynthetic growth (Paul *et al*., 1995; Boisset *et al*., 2023). The stabilisation of PRK by CP12 has been proposed in *A. thaliana* (López-Calcagno *et al*., 2017) and was supported by previous studies in *C. reinhardtii* (Gérard *et al*., 2022b; Teh *et al*., 2023). The previously proposed hypothesis of a direct molecular interaction between CP12 and PRK (Gérard *et al*., 2022b) in their reduced state may contribute to PRK stability and maintenance of RuBP levels under continuous light condition. There are few quantifications available of RuBP in Δ*CP12* lines or in *PRK* deficient lines. In the cyanobacterium *Synechocystis* sp. PCC 6803 under continuous light, deletion of the unique canonical *CP12* gene resulted in a greater amount of RuBP compared to the wild type upon a transition from high to low CO_2_ (Lucius *et al*., 2022). Under steady state conditions, the authors observed similar RuBP amounts in the WT and Δ*CP12* cells, but one hour after a transition from HC to VLC, the levels of RuBP increased more in the Δ*CP12* compared to the WT. In contrast, in *C. reinhardtii*, under steady state conditions, the lower the CP12 amount, the lower were PRK and RuBP amounts. We hypothesise that in eukaryotic microalgae, lower levels of RuBP trigger RuBisCO condensation in the pyrenoid, as does a low CO_2_/O_2_ ratio, and independently from other CCM components.

### Coordinated orchestration of RuBP recycling and CO_2_ acquisition by intrinsically disordered proteins

Altogether, our data showed that the absence of CP12 resulted in a lower amount of PRK and RuBP, and led to the absence of relocation of RuBisCO in the pyrenoid (Figure 5). We hypothesised that, in WT cells, the level of RuBP contributes to either the signalling or the adaptative response resulting in the relocation of RuBisCO to the pyrenoid, even though we cannot exclude that CP12 also contributes to this signalling alongside its effect on PRK and RuBP levels. Besides, these effects could be direct through molecular interaction or indirect (e.g., redox imbalance or phosphorylation cascade). In the absence of CP12, the distribution of RuBisCO in the stroma and the pyrenoid was independent of CO_2_ availability and resulted in a higher affinity for bicarbonate at HC. This alternative dissolved inorganic carbon acquisition is independent of the canonical CCM hallmarks and demonstrates a new interaction between RuBP recycling and CO_2_ acquisition.

**Fig. 5.**
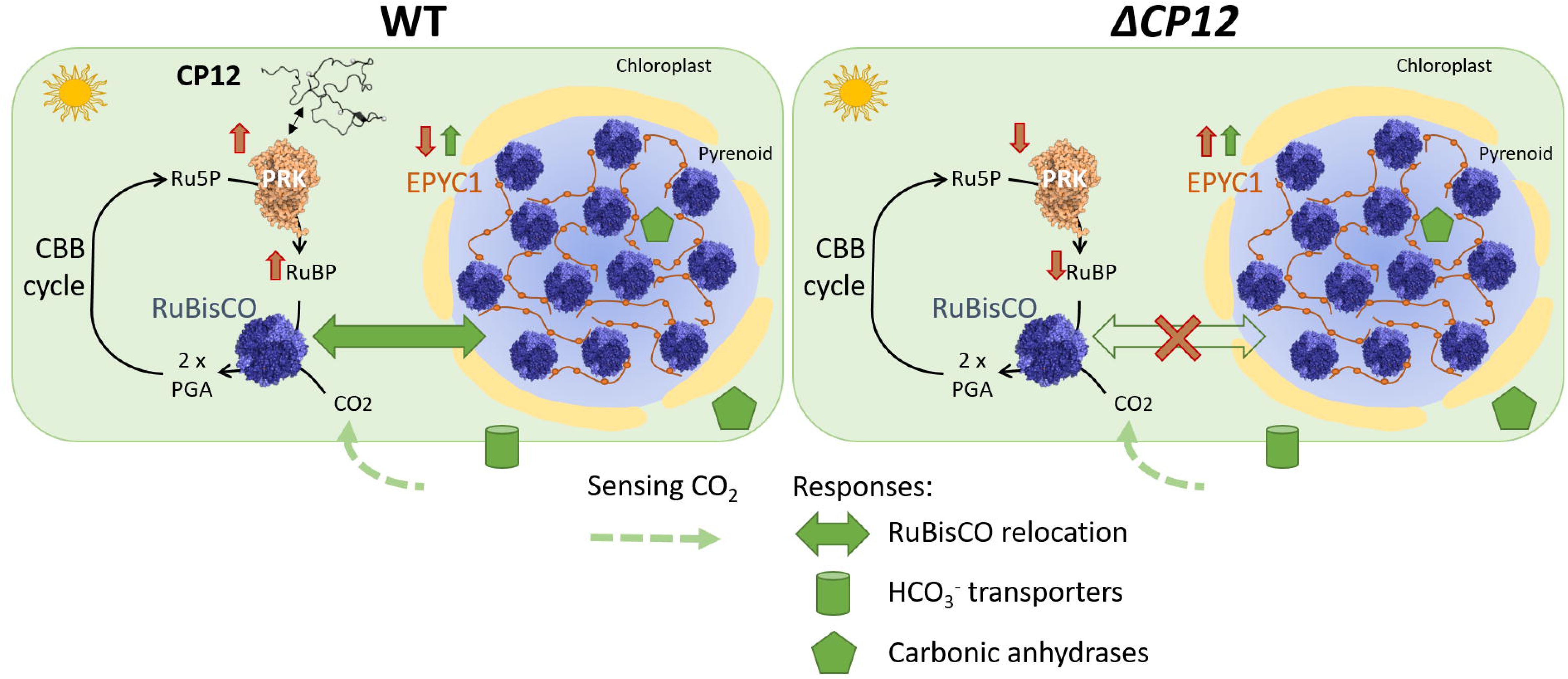
Schematic model of how CP12 absence may interfere with acclimation to VLC. The signal triggered by low CO_2_ concentration (green dashed arrow) is sensed and induces the expression of HCO_3_ ^-^ transporters (green cylinders), carbonic anhydrases (green hexagons) and the relocation of RuBisCO to the pyrenoid (green double arrow). The relocation of RuBisCO is associated with increased levels of EPYC1 (green simple arrow).The contribution of CP12 to this acclimation, revealed by comparing WT (left) and Δ*CP12* cells (right) is represented by the orange arrows and the orange cross. In Δ*CP12* cells, the level of PRK is reduced compared to WT cells, as well as the level of RuBP, and the relocation of RuBisCO is absent. The black double arrow indicates the molecular interaction between CP12 and PRK that could contribute to this effect (Gérard et al, 2022b). CBB: Calvin Benson Basham cycle (or reductive pentose phosphate pathway), Ru5P: Ribulose 5 phosphate, RuBP: Ribulose 1,5 bisphosphate, PRK: phosphoribulokinase, PGA: phosphoglycerate.

Interestingly, both CP12 and EPYC1 are intrinsically disordered proteins and are not uniformly distributed in the chloroplast and form punctate structures in *C. reinhardtii* (Wang *et al*., 2023) and in *Synechococcus elongatus* PCC 7942 (Perrin *et al*., 2025). The functions of the CP12 puncta are still enigmatic. In *Synechococcus*, a synergy between CP12, PRK and GAPDH condensation was observed. The relocation of the three proteins was induced by light, but was independent of the canonical dark-induced formation of the ternary complex (Perrin *et al*., 2025). This coordinated location of CP12 and PRK could be relevant regarding one of the functions we have identified here for CP12: the stabilisation of PRK and the maintenance of RuBP levels. The distribution of the CBB metabolic flux between these different punctae remains puzzling. Although the availability of CO_2_ in the pyrenoid has been extensively discussed, the availability of RuBP in the pyrenoid has been largely overlooked (Küken *et al*., 2018). Intrinsically disordered proteins are interesting candidates for orchestrating metabolic hubs, such as the CO_2_ acquisition and CO_2_ fixation that are both regulated by CP12.

## Acknowledgments

The technical service of the “Institut de la Microbiologie de la Méditerranée” is thanked for its valuable help: Jean-Michel Kaplansky, Alain Grossi, Evangelos Ailamakis and Gilles Kaczmarek. We thank Achille Marchand for help on the metabolite extraction protocol. We thank Xenie Johnson for providing the RuBisCO antibody raised against the large subunit shared by the Sprietzer laboratory. We also thank Pierre Crozet for providing the PRK antibody. We are grateful to Maya Belghazi for helpful discussions on proteomic data analysis.

## Author contributions

CG, RL, CV, HLG, AK, FG, KSC, LA were involved in acquisition, analysis and interpretation of data. CG, RL, LA, BG, EJ, SM, BGo and HL were involved in analysis and interpretation of data. KSC and EJ were involved in the design and the genetic construction of Δ*CP12* and Δ*CP12::CP12* strains; CG, LA, BGo and HL were involved in the experimental design, the acquisition and the primary analysis of the biochemical data. FG and BGa were involved in the experimental design, the acquisition and the primary analysis of the RuBP quantification. HLG and AK were involved in the experimental design of the electron microscopy experiments, the acquisition and the primary analysis of the images. CG, RL and CV were involved in the experimental design of the quantitative proteomic experiment, the acquisition and the primary analysis of proteomic data. CG was involved in the statistical analysis of the quantitative proteomic data. CG, SM, BGo and HL were involved in drafting the article and revising it critically for important intellectual content. BGo and HL were involved in the conception and design of the study, in analysis and interpretation of data, in drafting the article and revising it critically for important intellectual content. All authors were involved in the final approval of the version to be submitted.

## Supplementary materials

**Table S1** List of primers used for genomic PCR and quantitative real-time PCR in Cr*CP12* gene.

**Table S2** List of mass and charge of tryptic peptides of CP12 included in the MS/MS data acquisition programme for targeted quantification of CP12.

**Table S3** Summary of RuBisCO location by immunolabelling quantification.

**Table S4** List of accession numbers from Phytozome (*C. reinhardtii* CC-4532 v6.1) corresponding to the proteins highlighted in the volcano plots and heat maps in Fig.3.

**Table S5** Kinetics of photosynthesis of WT and Δ*CP12* strains grown at HC or VLC from fitting the data in Fig. 4 to equation 1 of the main text.

**Table S6** Kinetics of photosynthesis of WT and Δ*CP12* strains grown at HC or VLC from fitting the data in Fig. 4 to equation 2 of the main text.

**Fig. S1** Generation of Δ*CP12*::CP12, *CP12*-complemented transgenic strain (Δ*CP12*.compl.) and analysis of the genomic and transcriptional expression levels of *CP12*.

**Fig. S2** Analysis of GAPDH/CP12/PRK ternary complex formation by native electrophoresis and western-blot.

**Fig. S3** Western blot analysis of EPYC1 and PRK in WT and three independent mutants Δ*CP12* at HC and VLC. The abundance of CP12 at HC and VLC is also presented.

**Fig. S4** Phosphorylation state of EPYC1 at high CO_2_ (HC), very low CO_2_ VLC and in WT and Δ*CP12* strains.

**Fig. S5** Representative transmission electron microscopy (TEM) images of immunogold-labelled RuBisCO under different CO_2_ conditions.

**Fig. S6** Volcano plots displaying the changes of abundance of proteins in WT vs Δ*CP12* at HC or VLC.

**Fig. S7** Heatmap representing the abundance of proteins involved in carbon acquisition and assimilation in WT and Δ*CP12* at HC or VLC for each replicate.

**Fig. S8** Heatmap representing the abundance of all proteins in WT and Δ*CP12* (Δ) at high CO_2_ (HC) or at very low carbon (VLC) for each replicate.

## Data availability

**Supplementary data 1** Macro script used for analysis of the immunogold labelling images, also available here: https://github.com/Hugo-LE-GUENNO/macro-fiji/blob/main/PyreGold.ijm

**Supplementary data 2** Quantitative proteomic data (change in protein abundance on a log_2_ scale and statistical significance in –log₁₀ p-value) represented in the volcano plots Fig. 3A, 3B and S6.

**Supplementary data 3** Lists of proteins which abundance significantly changed upon transition from HC to VLC in WT and Δ*CP12* strains. Proteins common to both strains are highlighted in green.

**Supplementary data 4** Results of the ANOVA test for each replicate in WT and Δ*CP12* at HC or VLC (on a log2 scale). Proteins involved in carbon acquisition and assimilation represented in the heatmap Fig. S7 are in tab 2.

**Supplementary data 5** Results of Tukey’s Honest Significant Difference (THSD) test on the ANOVA significant hits. Proteins involved in carbon acquisition and assimilation represented in the heatmap Fig. S3D are in tab 2.

